# Frataxin Deficiency Drives a Shift from Mitochondrial Metabolism to Glucose Catabolism, Triggering an Inflammatory Phenotype in Microglia

**DOI:** 10.1101/2023.10.18.562916

**Authors:** Francesca Sciarretta, Fabio Zaccaria, Andrea Ninni, Veronica Ceci, Riccardo Turchi, Savina Apolloni, Martina Milani, Ilaria Della Valle, Marta Tiberi, Valerio Chiurchiù, Nadia D’Ambrosi, Silvia Pedretti, Nico Mitro, Katia Aquilano, Daniele Lettieri-Barbato

## Abstract

Immunometabolism investigates the complex interplay between the immune system and cellular metabolism. This study highlights the effects of mitochondrial frataxin (FXN) depletion, which causes Friedreich’s ataxia (FRDA), a neurodegenerative condition characterized by coordination and muscle control deficiencies. Using single-cell RNA sequencing, we identified specific cell groups in the cerebellum of a FRDA mouse model, emphasizing a notable inflammatory microglial response. These FXN-deficient microglia cells exhibited enhanced inflammatory reactions. Furthermore, our metabolomic analyses revealed increased glycolysis and itaconate production in these cells, possibly driving the inflammation. Remarkably, butyrate treatment counteracted these immunometabolic changes, triggered an antioxidant response via the itaconate-Nrf2-GSH pathways, and dampened inflammation. The study also pinpointed Hcar2 (GPR109A) as a potential agent for butyrate anti-inflammatory impact on microglia. Tests on FRDA mice highlighted the neuroprotective attributes of butyrate intake, bolstering neuromotor performance. In essence, our findings shed light on how cerebellar microglia activation contributes to FRDA and highlight butyrate potential to alleviate neuroinflammation, rectify metabolic imbalances, and boost neuromotor capabilities in FRDA and similar conditions.

## INTRODUCTION

Mutations in the frataxin (FXN) gene plays a critical role in the development of Friedreich’s ataxia (FRDA), a neurodegenerative disorder characterized by progressive muscle weakness and impaired coordination (Clark et al., 2018). In recent times, the significance of neuroinflammation in the context of neurodegenerative disorders has gained substantial recognition and emerging findings highlight a potential involvement of FXN loss in neuroinflammation (Apolloni et al., 2022). FXN deficiency has been shown to increase the production of pro-inflammatory cytokines, suggesting that FXN might be involved in regulating microglial activity (Hayashi et al., 2014; Khan et al., 2022; Shen et al., 2016). FXN is a mitochondrial protein, playing an essential role in the intricate process of iron-sulfur cluster assembly regulating mitochondrial electron transport chain (ETC) and aconitase activity. Loss of FXN has been suggested to disrupt mitochondrial oxidative capacity and cause mitochondrial ROS production (Al-Mahdawi et al., 2006; Anzovino et al., 2014). Aberrant mitochondrial metabolism and increased glycolytic flux are metabolic hallmarks of inflammatory macrophage/microglia activation (Jha et al., 2015; Sangineto et al., 2023). Although it is now well established that FXN takes center place in mitochondrial metabolism, the consequence of FXN loss in microglia cells has never been explored. Several authors demonstrated that the mitochondrial metabolite itaconate causes Krebs cycle break and modulates inflammatory response. Itaconate inhibits glycolysis flux and oxidative stress, limiting the inflammatory setting in macrophages and microglia (Lampropoulou et al., 2016; Pan et al., 2023). Similarly, forcing mitochondrial oxidative metabolism improves the inflammatory phenotype in macrophages. It has been reported that microbiota-derived short-chain fatty acid (SCFA) butyrate enhances oxidative metabolism and uncouples Krebs cycle from glycolytic flux in immune cells (Bachem et al., 2019). Butyrate had a neuroprotective impact on mouse models of Parkinson’s disease, likely due to the downstream regulation of gut microbiota and inhibition of gut-brain axis inflammation (Guo et al., 2023). Butyrate reduces neuroinflammation and microglia activation in several experimental models of disease (Caetano-Silva et al., 2023; Huuskonen et al., 2004; Wenzel et al., 2020). Butyrate has been identified as a high-affinity ligand for the Gi-linked heterotrimeric guanine nucleotide-binding protein–coupled receptor (GPCR) hydroxycarboxylic acid receptor 2 (HCAR2) (Carretta et al., 2021), which is expressed in the brain and has been shown to modulate microglial actions in several neuroinflammatory diseases such as multiple sclerosis, Parkinson’s disease, Alzheimer’s disease (Moutinho et al., 2022; Offermanns, 2014). Recently, diminished abundance of butyrate-producing bacteria has been demonstrated in a mouse model of FRDA (Turchi et al., 2023). Dietary butyrate supplementation in FRDA mice limited macrophage activation in white adipose tissue and in bone marrow-derived macrophages, suggesting that this molecule could be also efficient in mitigating neuroinflammation and neurobehavior disability. Herein we demonstrated the FXN loss causes an immunometabolic derangement in microglial cells enhancing glucose catabolism to sustain a strong inflammatory phenotype. This evidence was corroborated in an *in vivo* model of FRDA. Butyrate effectively restored the immunometabolic defects both in vitro and in vivo improving the neuromotor abilities in the FRDA mouse model.

## RESULTS

### Frataxin deficiency activates cerebellar microglia of KIKO mice

Although genetic deficiency of the mitochondrial protein FXN is causative of FRDA-related symptoms, the disease-specific cell types in cerebellum are unknown yet. Herein we used a comparative single cell RNA-sequencing (scRNA-seq) between cerebellum of WT and KIKO mice at the early stage of FRDA disease (6-months old). ScRNA-seq analysis led to the identification of a total of 4484 quality control (QC)-positive cells. Based on the expression levels of the most variable genes, we annotated homogeneous and robust cluster of cells from scRNA-seq data, resulting in 11 group of cells such as granule cells (GC), T cells, oligodendrocytes, NK cells, microglia/macrophages, fibroblasts, ependymal cells, external granular layer cells (EGL), choroid plexus cells, blood cells, astrocytes (**Fig. 1A** and **Fig. 1B**). To avoid subjectivity and to add strength to the analyses, we also performed reference-based single-cell annotation, which confirmed our broad clusters of cell populations. Next, by using cell-marker gene, we compared the cell clusters between genotype and we found a contraction of choroid plexus cells, astrocytes and oligodendrocytes (**Fig. 1C**). Oppositely, increased markers of microglia/macrophage population in cerebellum of KIKO were observed (**Fig. 1B** and **Supplemental Fig. 1A**). Consistently, the gene ontology (GO) terms for biological processes revealed an enrichment in the inflammatory process and downregulation of mitochondrial oxidative genes (**Fig. 1D**).

**FIGURE 1.**
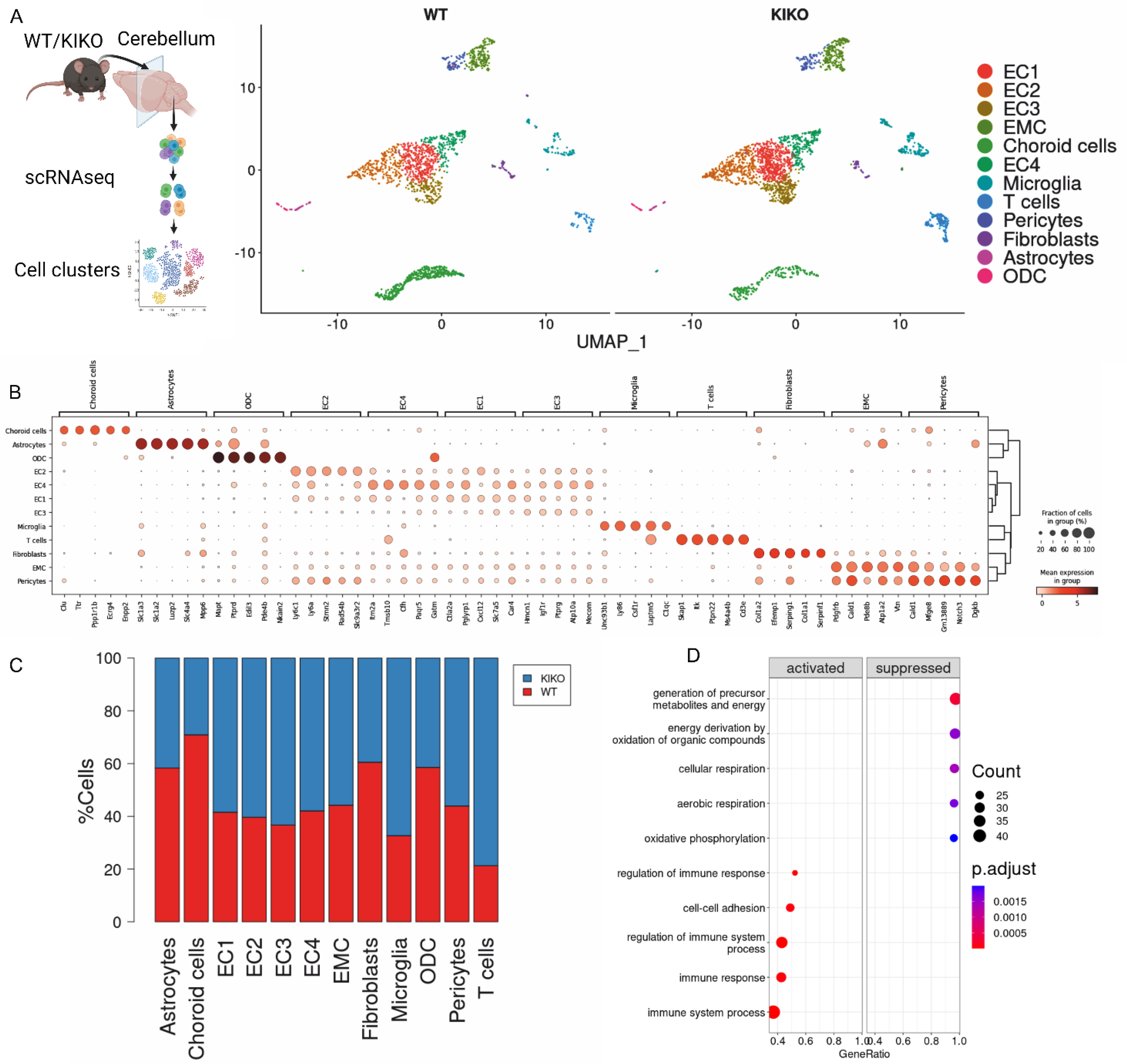
Cerebellum in KIKO exhibits an immunometabolic disturbance. (**A**) Cell clusters identified by single cell RNA-seq of total cell fraction (TCF) isolated from cerebellum of WT and KIKO of 6 months old mice (TCF: pool from cerebellum of n=4 mice/group). (**B**) Dot plots reporting gene markers for cell type identified by single cell RNA-seq of total cell fraction (TCF) isolated from cerebellum of WT and KIKO of 6 months old mice (TCF: pool from cerebellum of n=4 mice/group). (**C**) Bar plots reporting cell types identified by single cell RNA-seq of total cell fraction (TCF) isolated from cerebellum of WT and KIKO of 6 months old mice (TCF: pool from cerebellum of n=4 mice/group) (**D**). Gene Ontology terms for biological processes of differentially expressed genes revelated by single cell RNA-seq of total cell fraction (TCF) isolated from cerebellum of WT and KIKO of 6 months old mice (TCF: pool from cerebellum of n=4 mice/group).

Microglia are the primary innate immune cells of the central nervous system (CNS) that are sentinels participating in the inflammatory and cell clearance response (Norris and Kipnis, 2019). To corroborate the microglia dynamics of KIKO mice, we performed a high dimensional flow cytometry analysis (**Fig. 2A**) and we observed that although the percentage of cerebellar CD45^low^CD11b+ microglia remained preserved, a higher expression of M1-like markers (CD86^+^ and MHC-II) and a concomitant lower expression of M2-like marker CD206 was observed in microglial cells of KIKO compared to WT (**Fig. 2B**). No changes were observed in the percentages of neutrophils (CD45+/CD11b-/CD3-/NK1.1+/CD90.2-cells) and B cells (CD45+/CD11b-/BB220+/Ly6G+ cells) (**Supplemental Fig. 1B**), whereas a significant increase was detected in T cells (CD45+/CD11b-/BB220-/Ly6G-/CD3+/CD90.2+ cells) (**Supplemental Fig. 1C**). Next, to explore the molecular signatures of microglia, cerebella CD45^+^/CD11b^+^ cells were isolated by magnetic cell sorting and their transcriptome was profiled by bulk RNA-sequencing. We identified n=1275 differentially expressed genes (−0.75>Log2FC>+0.75; p<0.05) between cerebella microglia of KIKO vs WT mice (**Fig. 2C**). GO terms for biological processes of the top 200 down-regulated genes (orange bars) revealed a reduced mitochondrial oxidative capacity in microglia/macrophages of KIKO mice (**Fig. 2D**); in an opposite manner, the top 200 up-regulated genes (green bars) pertained to inflammatory processes as well as response to inflammatory stimuli (**Fig. 2E**). These results suggest a limited mitochondrial oxidative capacity with an increased inflammatory phenotype in microglia of KIKO mice.

**FIGURE 2.**
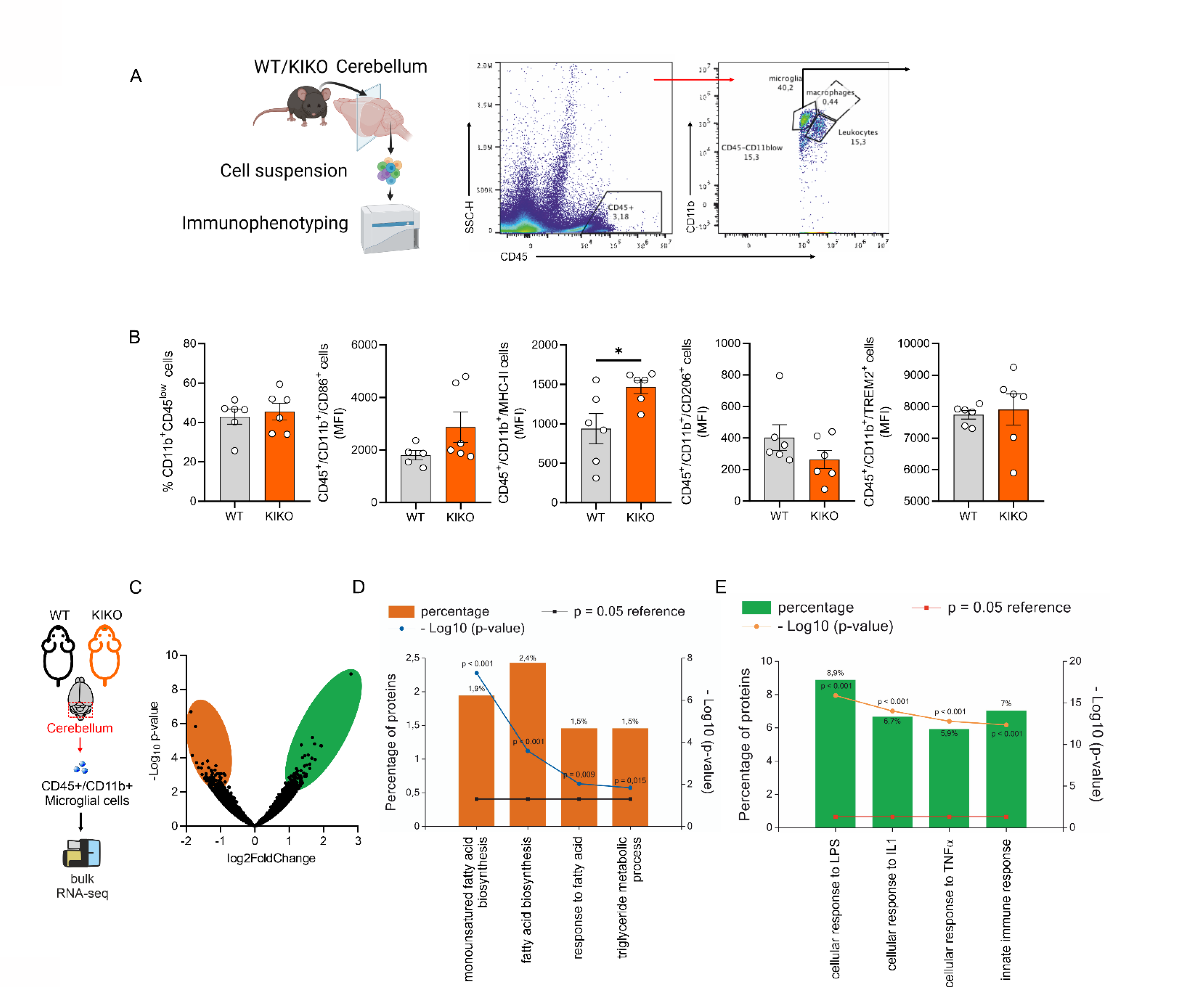
Microglia-derived from cerebellum of KIKO mice shows an inflammatory phenotype. (**A, B**) High dimensional flow cytometry of pro-inflammatory (CD86 and MHC-II) and anti-inflammatory (CD206 and Trem2) markers in microglial cells (CD4^low^+/CD11b^+^) isolated from cerebellum of WT and KIKO of 6 months old mice (n=5/6 mice/group). Data were reported as mean ± SD. Student’s t test ^*^ p<0.05. (**C**) Volcano plot of differentially expressed genes (DEGs: -0.75>Log2FC>+0.75; p<0.05) in microglia isolated from 6 months old KIKO and wild type (WT) mice (n=4 mice/group). (**D, E**) Functional enrichment analysis for biological processes of downregulated genes (D, orange bars) and up-regulated genes (E, green bars) in microglia isolated from 6 months old KIKO and wild type (WT) mice (n=4 mice/group).

### Loss of frataxin enhance glucose catabolism in microglial cells

To give more insight to the molecular mechanisms leading to inflammatory activation observed in KIKO-derived microglia cells, we generated a FRDA cell model by stably downregulating FXN in a microglia cell line (BV2^FXN-^). Although, the analysis of the inflammatory profile did not reveal any difference between controls (BV2^SCR^) and BV2^FXN-^ cells (**Fig. 3A**), under basal conditions, the highest inflammatory susceptibility was observed when BV2^FXN-^ cells were activated with liposaccharides (LPS) (**Fig. 3A**). These results suggest that activated BV2^FXN-^ better phenocopy the inflammatory setting observed in cerebella microglia of KIKO mice.

**FIGURE 3.**
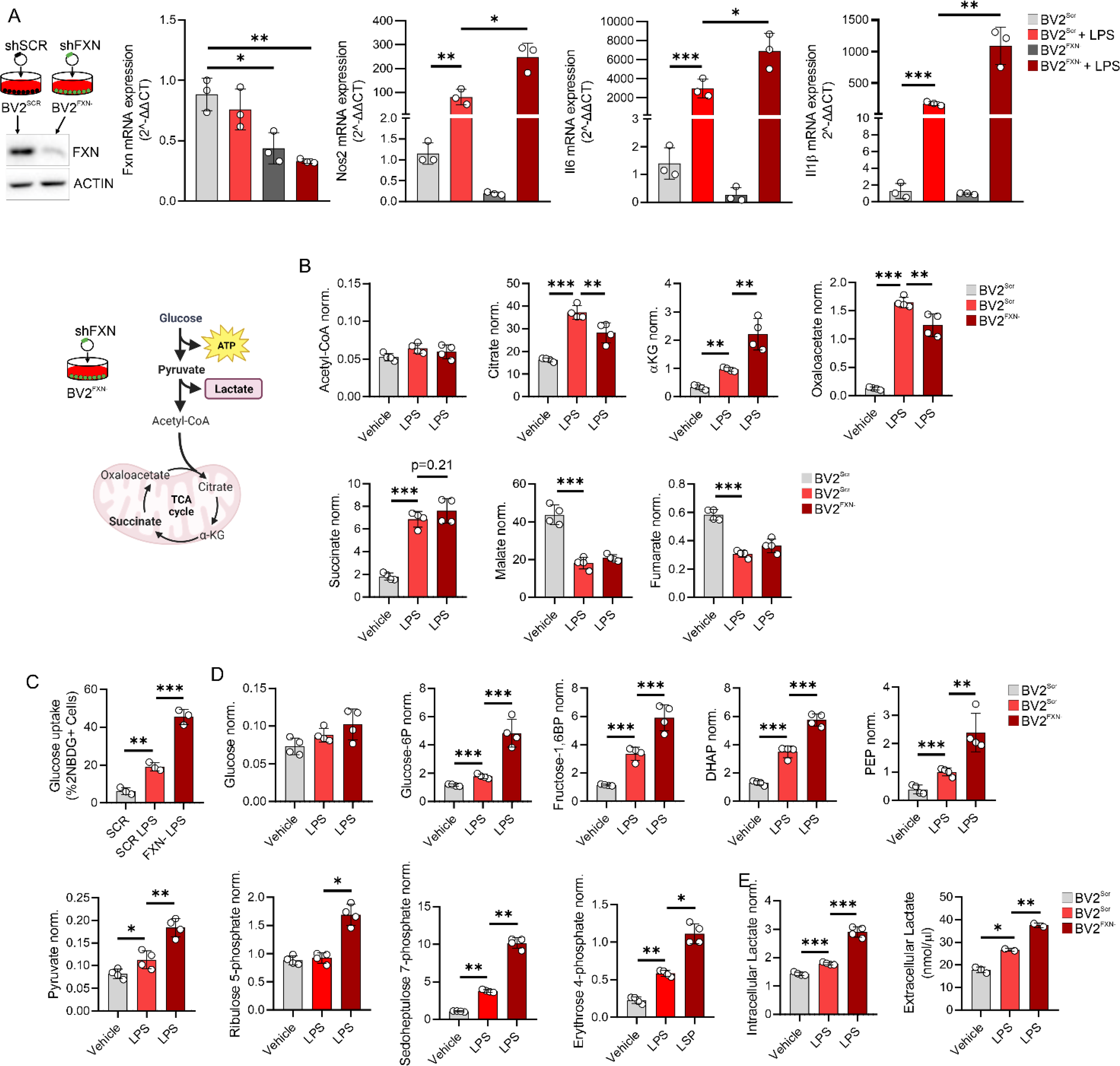
Loss of frataxin forces glucose catabolism in microglial cells. (**A**) BV2 cells were transfected with lentiviral particles delivering Fxn or Scr sequence and gene expression level of inflammatory genes (Nos2, Il6, Il1β) were analyzed by qPCR. LPS (500 ng/mL for 16 hours) was used to activate BV2 cells. Data were reported as mean ± SD. ANOVA test ^*^ p<0.05; ^**^ p<0.01; ^***^ p<0.001. (**B**) BV2 were cells transfected with lentiviral particles delivering Fxn or Scr sequence and metabolites tracking TCA cycle were measured by LC-MS. LPS (500 ng/mL for 16 hours) was used to activate BV2 cells. Data were reported as mean ± SD. ANOVA test ^**^ p<0.01; ^***^ p<0.001. (**C**) BV2 cells transfected with lentiviral particles delivering Fxn or Scr sequence were loaded 2NBDG for 30 minutes. Glucose uptake calculated as 2-NBDG+ cells by flow cytometry. LPS (500 ng/mL for 16 hours) was used to activate BV2 cells. Data were reported as mean ± SD. ANOVA test ^**^ p<0.01; ^***^ p<0.001. (**D**) BV2 were cells transfected with lentiviral particles delivering Fxn or Scr sequence and metabolites tracking glycolysis and pentose phosphate pathway were measured by LC-MS. LPS (500 ng/mL for 16 hours) was used to activate BV2 cells. Data were reported as mean ± SD. ANOVA test ^*^ p<0.05; ^**^ p<0.01; ^***^ p<0.001. (**E**) BV2 were cells transfected with lentiviral particles delivering Fxn or Scr sequence and lactate production was measured by LC-MS (intracellular) or spectrophotometer (extracellular). LPS (500 ng/mL for 16 hours) was used to activate BV2 cells. Data were reported as mean ± SD. ANOVA test ^*^ p<0.05; ^**^ p<0.01; ^***^ p<0.001.

It has been demonstrated that a sustained inflammatory status of macrophages is characterized by metabolic shift from oxidative phosphorylation (OXPHOS) to glycolysis. To test if a metabolic rearrangement occurs in FXN downregulating microglia, we measured glycolysis- and TCA-related metabolites in BV2^FXN-^ cells. Notably, mitochondrial metabolites including acetyl-CoA, citrate, α-ketoglutarate (αKG), oxaloacetate, succinate and malate were unchanged in activated BV2^FXN-^ cells (**Fig. 3B**). On the contrary, glucose uptake (**Fig. 3C**), accumulation of glycolysis and pentose-phosphate shunt metabolites (**Fig. 3D**) as well as lactate production (**Fig. 3E**) were significantly increased. These results were consistent with the glucose avidity of inflammatory immune cells (Soto-Heredero et al., 2020). With the aim to explore if the inflammatory phenotype of BV2^FXN-^ cells was dependent on glycolysis, we inhibited glucose uptake by 2-deoxyglucose (2-DG) and as reported in the **Suppl. Fig. 2A**, a reduced inflammatory response to LPS was observed.

### Itaconate reduces the inflammatory responses in microglia downregulating FXN through Nrf2 pathway

Itaconate is a mitochondrial metabolite produced in macrophages as response to inflammatory stimuli (Lampropoulou *et al*., 2016). To test if the highest inflammatory phenotype of BV2^FXN-^ cells was also associated with itaconate overproduction, we measured its levels, and a significant increase was detected compared to scramble conditions (with or without LPS) (**Fig. 4A**). Itaconate overproduction observed in BV2^FXN-^ cells was in accordance with the increased expression levels the immune-responsive gene 1 (Irg1) (**Fig. 4B**), the mitochondrial enzyme catalyzing the decarboxylation of cis-aconitate to synthesize itaconate (Lampropoulou *et al*., 2016). Remarkably, higher Irg1 expression levels were also detected in cerebellum-derived microglia (**Fig. 4C**) as well as in total cerebellum of KIKO than WT mice (**Fig. 4D**). It has been reported that itaconate production following inflammatory stimuli mitigates the inflammatory response by constraining Il1b induction and glycolysis (Lampropoulou *et al*., 2016). To test if itaconate exerts such effect in activated BV2^FXN-^ cells, a cell permeable formulation of soluble itaconate (dimethylitaconate: DMI) was added to the culture medium. As expected, DMI diminished Il1β, Il6 and Nos2 levels (**Fig. 4E**) as well as glucose catabolism in BV2^FXN-^ cells (**Fig. 4F**). It has been demonstrated that itaconate exerts its anti-inflammatory effect by Nrf2 induction (Mills et al., 2018).

**FIGURE 4.**
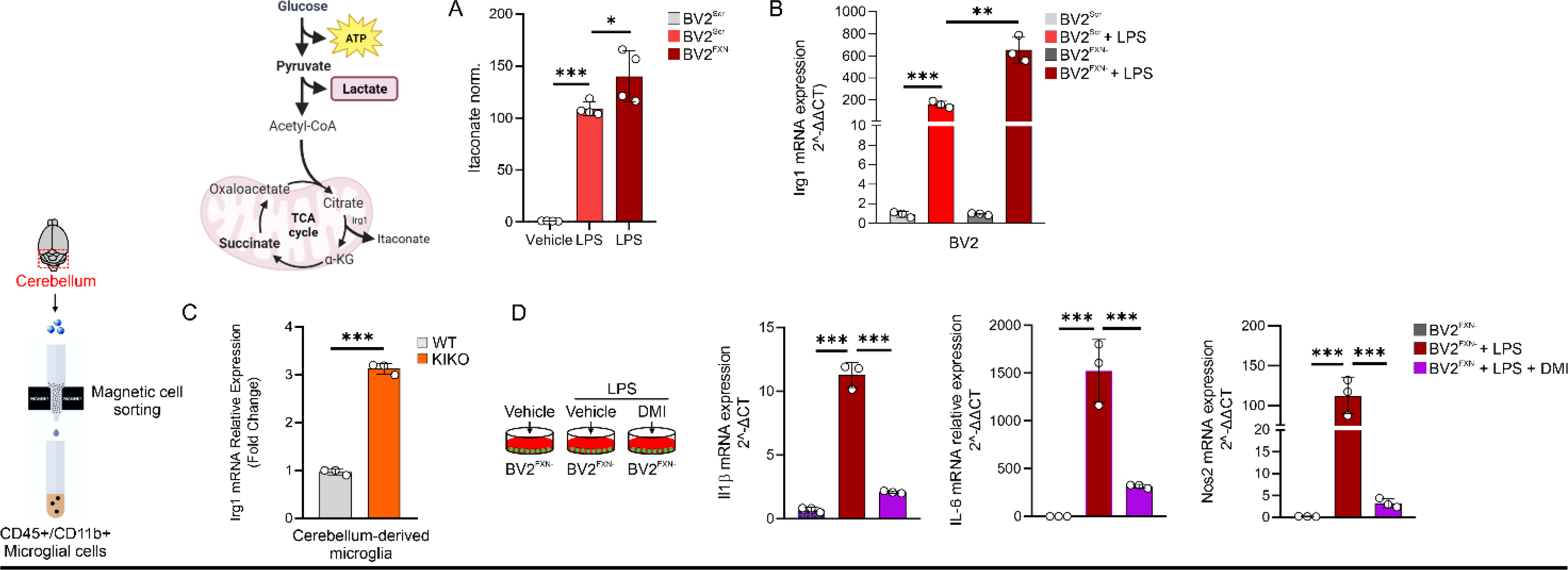
Itaconate overproduction restrains the inflammatory phenotype in FRDA microglial cells. (**A, B**) BV2 cells were transfected with lentiviral particles delivering Fxn or Scr sequence and itaconate production (A) and Irg1 mRNA expression (B) were analyzed by LC-MS and qPCR, respectively. LPS (500 ng/mL for 16 hours) was used to activate BV2 cells. Data were reported as mean ± SD. ANOVA test ^*^ p<0.05; ^**^ p<0.01; ^***^ p<0.001. (**C**) Microglial cells were isolated from cerebellum of 6 months old KIKO or WT mice by magnetic cell sorting (CD45^+^/CD11b^+^ cells) and Irg1 mRNA expression was analyzed by qPCR. Data were reported as mean ± SD. Student’s t test ^***^ p<0.001. (**D**) BV2 cells were transfected with lentiviral particles delivering Fxn sequence and the inflammatory gene expression was analyzed by qPCR. LPS (500 ng/mL for 16 hours) was used to activate BV2 cells. Dimethyl itaconate (DMI, 100M) was added 3 hours before LPS treatment and maintained throughout the experiment. Data were reported as mean ± SD. ANOVA test ^***^ p<0.001).

### Butyrate reverts the immunometabolic signatures through Itaconate/Nrf2/GSH signaling

Butyrate is a ubiquitous short-chain fatty acid principally derived from the enteric microbiome, which showed a neuroprotective role (Lanza et al., 2019; Li et al., 2016). Metabolomic analysis of butyrate-treated macrophages revealed a substantial reduction in glycolysis (Flemming, 2019; Schulthess et al., 2019) as well as limited inflammatory response in microglia (Caetano-Silva *et al*., 2023). By virtue of the recently demonstrated anti-inflammatory effects of BUT on white adipocytes and BMDM of KIKO mice (Turchi *et al*., 2023), we asked if butyrate treatment was also effective in counteracting the changes of the immunometabolic profile in activated BV2^FXN-^. As reported in **Fig. 5A**, butyrate reduced glucose uptake and lowered lactate production in activated BV2^FXN-^ cells, whereas a significant refill in the mitochondrial metabolites such as citrate, oxaloacetate and succinate was observed (**Fig. 5B**). Notably, butyrate further increased itaconate levels in BV2^FXN-^ (**Fig. 5C**), leading us to suppose that butyrate promotes an anti-inflammatory effect by the itaconate-driven antioxidant protection. To test this hypothesis, we analyzed Nrf2 protein in BV2^FXN-^ cells treated with butyrate and expectedly an increased nuclear accumulation of Nrf2 was observed (**Fig. 5D**). Nrf2 is the primary transcription factor protecting cells from oxidative stress by regulating the synthesis of glutathione (GSH) (Harvey et al., 2009). Interestingly, butyrate increased GSH levels in BV2^FXN-^ cells (**Fig. 5E**).

**FIGURE 5.**
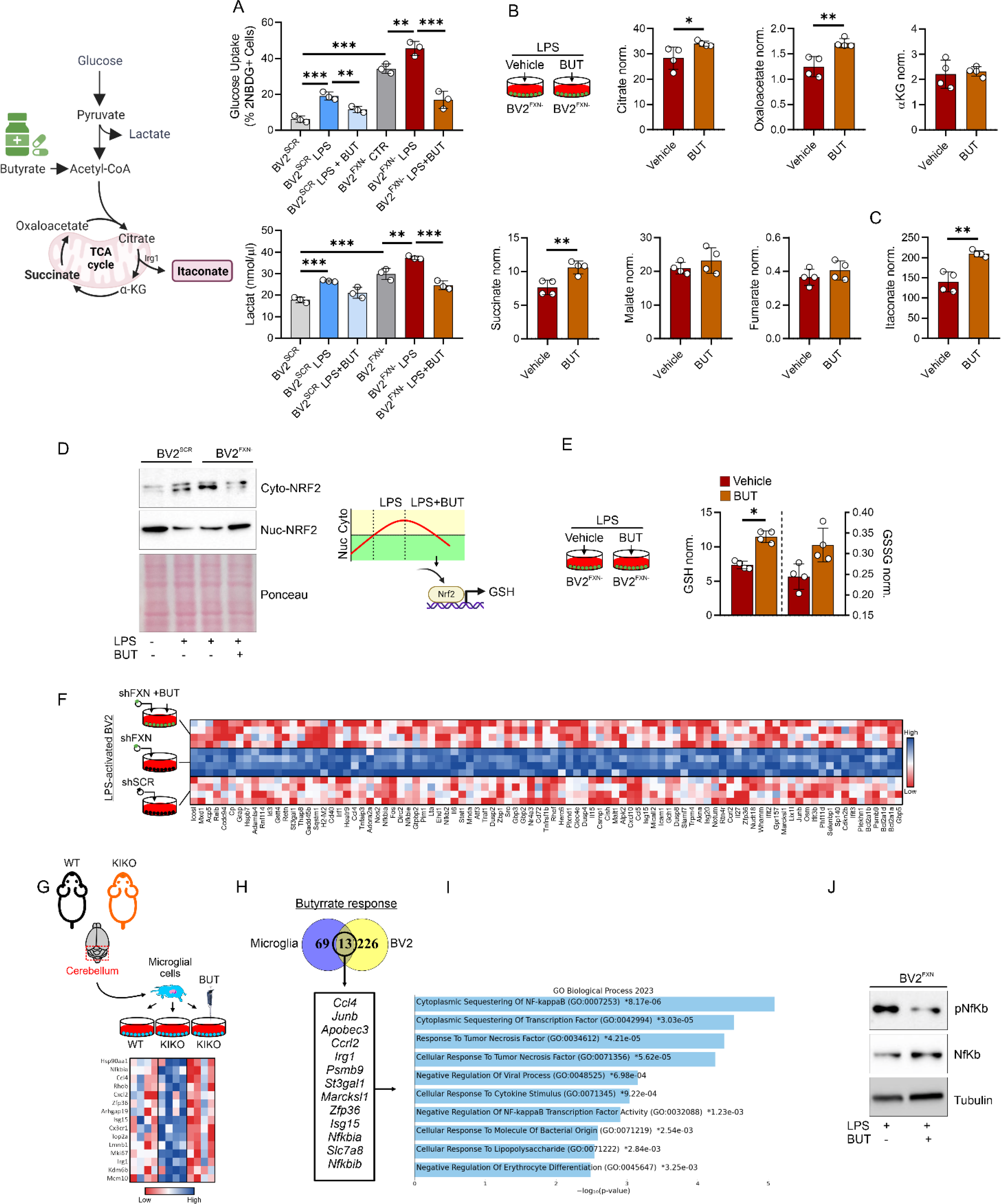
Butyrate rewires the immunometabolism of microglia downregulating FXN. (**A**) BV2 cells were transfected with lentiviral particles delivering Fxn or Scr sequence and glucose uptake (upper panel) and lactate production (bottom panel) were measured by flow cytometry and spectrofluorometer, respectively. LPS (500 ng/mL for 16 hours) was used to activate BV2 cells. Sodium butyrate (BUT, 500μM) was added 3 hours before LPS treatment and maintained throughout the experiment. Data were reported as mean ± SD. ANOVA test ^**^p<0.01; ^***^ p<0.001. (**B, C**) BV2 cells were transfected with lentiviral particles delivering Fxn sequence and metabolites tracking TCA cycle (B) and itaconate (C) were measured by LC-MS. LPS (500 ng/mL for 16 hours) was used to activate BV2 cells. Sodium butyrate (BUT, 500μM) was added 3 hours before LPS treatment and maintained throughout the experiment. Data were reported as mean ± SD. Student’s t test ^*^ p<0.05; ^**^p<0.01). (**D**) BV2 cells were transfected with lentiviral particles delivering Fxn or Scr sequence and cytosolic/nuclear fractions of NRF2 were analyzed by western blot. LPS (500 ng/mL for 16 hours) was used to activate BV2 cells. Sodium butyrate (BUT, 500μM) was added 3 hours before LPS treatment and maintained throughout the experiment. Ponceau staining was used as loading control. (**E**) BV2 cells were transfected with lentiviral particles delivering Fxn sequence and GSH and GSSG levels were measured by LC-MS. LPS (500 ng/mL for 16 hours) was used to activate BV2 cells. Sodium butyrate (BUT, 500μM) was added 3 hours before LPS treatment and maintained throughout the experiment. Data were reported as mean ± SD. Student’s t test ^*^ p<0.05. (**F**) Heatmap of differentially expressed genes (p<0.05) in BV2 cells transfected with lentiviral particles delivering Fxn or scramble (Scr) sequence. LPS (500 ng/mL for 16 hours) was used to activate BV2 cells. Sodium butyrate (BUT, 500μM) was added 3 hours before LPS treatment and maintained throughout the experiment. (**G**) Heatmap of differentially expressed genes (p<0.05) in microglia isolated from the cerebellum of WT and KIKO mice. Sodium butyrate (BUT, 500μM) was added to the culture medium for 16 hours. (**H, I**) Venn diagram of butyrate-responsive genes in LPS-stimulated BV2 and microglia isolated from KIKO mice (H) and the functional enrichment analysis of the overlapping genes was analyzed by EnrichR (I). (**J**) BV2 cells were transfected with lentiviral particles delivering Fxn sequence and pospho-active and basal form of NfKb were analyzed by western blot. Tubulin was used as loading control. LPS (500 ng/mL for 16 hours) was used to activate BV2 cells. Sodium butyrate (BUT, 500μM) was added 3 hours before LPS treatment and maintained throughout the experiment.

In order to decipher the molecular mechanism driving the anti-inflammatory effect of butyrate, we analyzed the transcriptomics responses to butyrate in BV2^FXN-^ (**Fig. 5F)** and KIKO-derived microglial cells **(Fig. 5G**). The genes that were significantly downregulated (Log2FC<-1.5) by butyrate were integrated by Venn diagram (**Fig. 5H**) and their functional enrichment analysis suggested that butyrate inhibits NfkB signaling pathway in microglia with FXN deficiency (**Fig. 5I**). Consistently, a diminished level of the phospho-active form of Nf-κb was observed in activated BV2^FXN-^ treated with butyrate (**Fig. 5J**). To demonstrate the anti-inflammatory effects of butyrate *in vivo*, asymptomatic 4-months-old KIKO mice were fed with dietary BUT for 16 weeks and at the end of dietary treatment, the transcriptome of CD11b^+^ microglial cells isolated from cerebellum, was profiled. In accordance with *in vitro* data, KIKO-derived CD11b^+^microglial cells showed a reduced expression level of inflammatory genes following dietary BUT treatment (**Suppl. Fig. 2B**).

### Butyrate improves the neuromotor abilities in KIKO mice

Next, we asked if the improvement of the neuroinflammatory status of BUT-treated mice was accompanied by improved neuromotor abilities. To this end, a battery of neuromotor tasks including accelerating rotarod test (Bohlen et al., 2009), pole tests (turning time and climb down time) (Que et al., 2021) and tightrope test (Miquel and Blasco, 1978) were conducted in KIKO mice at the end of dietary treatment. The rotarod test revealed lower neuromotor capacity in KIKO mice compared to the WT mice when the mice ran at maximum RPM (**Fig. 6A**). Nicely, butyrate treatment was effective in limiting KIKO falls (**Fig. 6A**). Similar results were observed following pole test, in which KIKO mice showed a highest time to turn completely downward (Tturn) and to descend to the floor (Ttotal) than WT mice (**Fig. 6B**). Although butyrate treatment was effective in improving Tturn (**Fig. 6B**), no improvement was observed in Ttotal (**Fig. 6B**). Restored neurobehavioral abilities were also observed at the tightrope test, in which butyrate reduced the higher walking time of KIKO than WT mice (**Fig. 6C**). These results suggest that dietary butyrate improves neuromotor abilities through neuroinflammatory limitation in FRDA mice.

**FIGURE 6.**
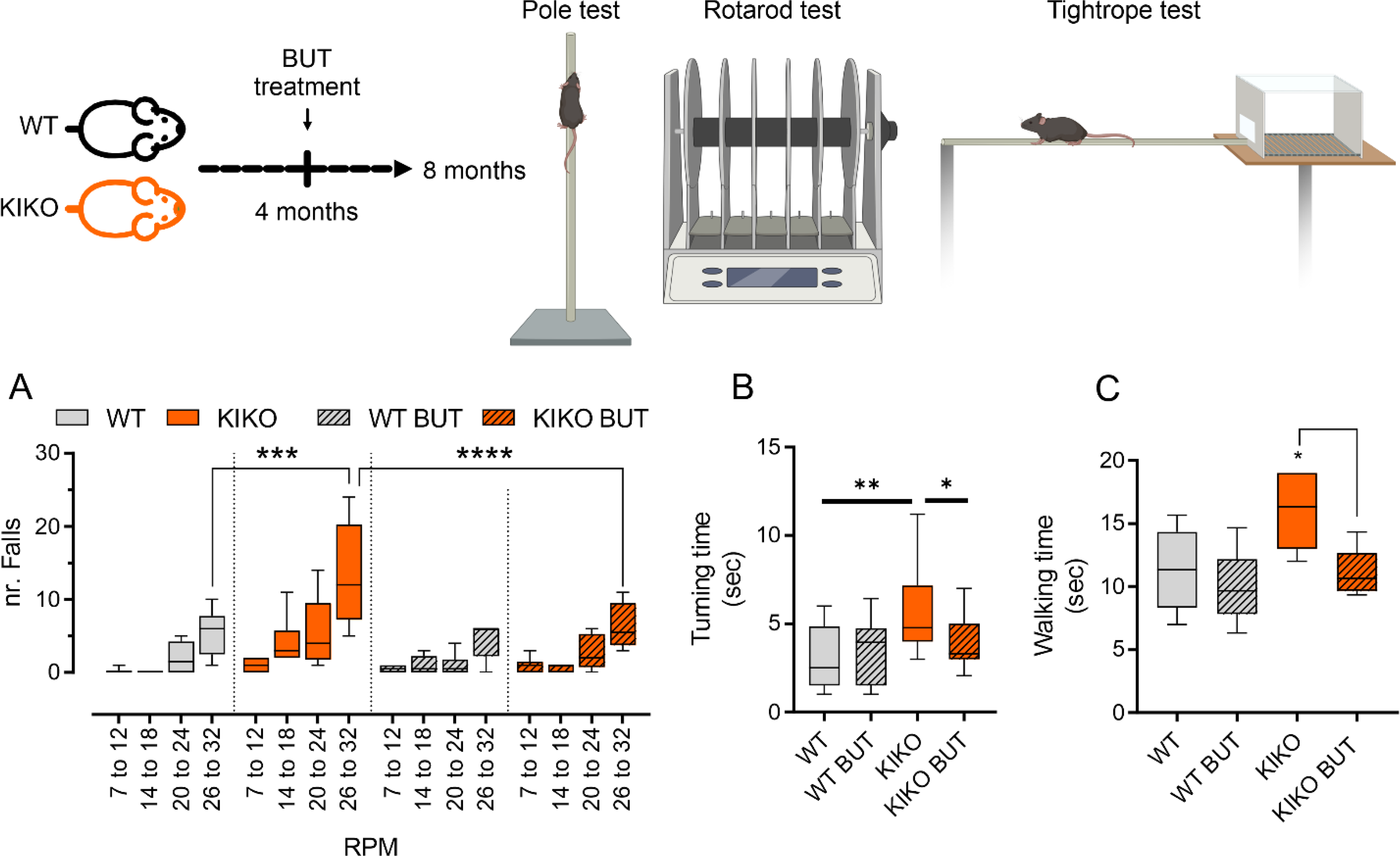
Butyrate supplementation enhances neuromotor performance in KIKO mice. Male WT and KIKO mice, aged four months, were either maintained on a standard diet or one supplemented with butyrate (BUT) for a duration of 16 weeks, until they reached eight months of age. **A)** Rotarod test performance, expressed by the number of falls, across various speeds. B) Duration taken for the mice to turn during pole test atop the pole. **C**) Time of walking during the tightrope test. Data are presented as mean ± SD. ANOVA ^*^ p<0.05, ^**^ p<0.01, ^***^ p<0.001, ^****^ p<0.0001 (n=6 mice/group).

### Hcar2 mediates the anti-inflammatory effects of butyrate

Butyrate interacts with several G-protein coupled receptors including GPR109A (encoded by Hcar2 gene), GPR43 (encoded by Ffar2 gene) and GPR41 (encoded by Ffar3 gene) leading to activation of anti-inflammatory signaling cascades (Deleu et al., 2021; Parada Venegas et al., 2019). Through According to what previously reported (Moutinho *et al*., 2022), our scRNAseq data revealed that among these receptors Hcar2 was expressed at the highest values in microglia (**Fig. 7A)**. Interestingly, the Hcar2 expression was higher in in KIKO than WT mice (**Fig. 7B)**, suggesting an increased sensitivity to butyrate in FRDA mice. Remarkably, Hcar2 levels were increased in primary microglia isolated from cerebellum of KIKO mice (**Fig. 7C**), and butyrate treatment was effective in restraining its upregulation (**Fig. 7C**). In line with these findings, butyrate limited the expression levels of Hcar2 in activated BV2^FXN-^ cells (**Fig. 7D**). To investigate if Hcar2 mediates the anti-inflammatory effects of butyrate, we downregulated Hcar2 in butyrate-pre-treated BV2^FXN-^ cells and the inflammatory genes expression was analyzed following LPS stimulation. Consistent with our hypothesis, butyrate was ineffective in limiting inflammatory response in Hcar2 downregulating cells (**Fig. 7E**).

**FIGURE 7.**
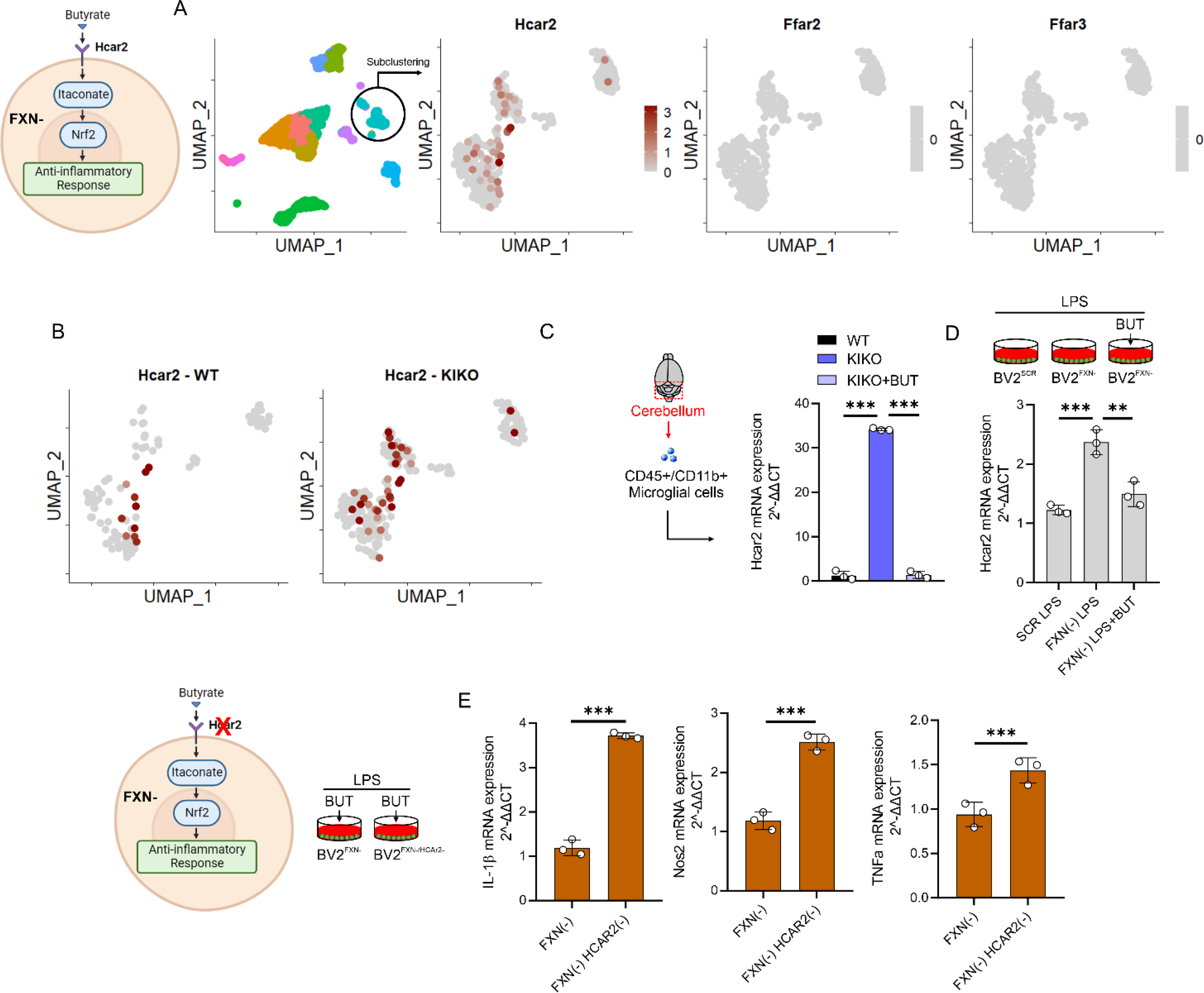
Hcar2 mediates the butyrate responses in the FRDA microglia. (**A**) Cerebellar microglial cells analyzed by scRNA-seq were subclustered and Hcar2, Ffar2 and Ffar3 expression levels were analyzed (pool of n=4 mice/group). (**B**) Hcar2 expression levels in cerebellar microglia of 6 months-old WT and KIKO mice (pool of n=4 mice/group). (**C**) Microglia were isolate from cerebellum of WT, KIKO or KIKO mice fed butyrate and Hcar2 expression level was measured by qPCR (n=3 mice/group. Data were reported as mean ± SD. ANOVA test ^***^p<0.001). (**D**) BV2 cells were transfected with lentiviral particles delivering scramble (Scr) or Fxn sequence and Hcar2 expression levels and Hcar2 expression level was measured by qPCR. Data were reported as mean ± SD. ANOVA test ^**^p<0.01; ^***^p<0.001.

## DISCUSSION

FXN deficiency caused excess microglial DNA damage and inflammation in murine model of FRDA (Shen *et al*., 2016). Remarkably, the transcriptional profile of PBMC isolated from FRDA patients, revealed a strong enrichment for an inflammatory innate immune response (Nachun et al., 2018). Although high inflammatory susceptibility was described in FRDA mice and human (Khan *et al*., 2022; Shen *et al*., 2016; Turchi *et al*., 2023), the mechanisms underlying this condition remain unexplored. Herein we demonstrated that the loss of FXN forces glycolytic catabolism promoting inflammatory phenotype in microglial cells. FXN is a mitochondrial protein and its dysfunction causes mitochondrial failure, thus recruiting glycolysis as the main source of ATP (O’Neill et al., 2016). This is consistent with the increased glycolysis flux occurring in M1 macrophages and microglial cells (Bernier et al., 2020). Krebs cycle breaks were also described in M1 macrophages and microglia, which cause an overproduction of itaconic acid (Lampropoulou *et al*., 2016). This mitochondrial metabolite has been shown to participate in the inflammatory response restraining IL1β production and glycolysis. Itaconate and its derivatives showed antiinflammatory effects in preclinical models of sepsis, viral infections, psoriasis, gout, ischemia/reperfusion injury, and pulmonary fibrosis, pointing to possible itaconate-based therapeutics for a range of inflammatory diseases (Peace and O’Neill, 2022). Consistently, itaconate improved the immunometabolic profile in microglia downregulating FXN through Nrf2-mediated mechanism, highlighting itaconate as novel therapeutical option to improve FRDA-related inflammatory symptoms. It has been reported that itaconate exerts its anti-inflammatory role by activating Nrf2 (Mills *et al*., 2018). Nrf2 controls the antioxidant responses counteracting the production of oxidatively damaged molecules through GSH synthesis (Mills *et al*., 2018). Of note, Nrf2 is down-regulated in FRDA patients and antioxidant GSH precursors improve FRDA symptoms (La Rosa et al., 2021).

Mounting evidence reports that gut microbiota releases immunomodulatory molecules and counteracts neuroinflammatory conditions. (Abdel-Haq et al., 2019; Mou et al., 2022; Richards et al., 2022; Sampson et al., 2016). To this end targeting gut microbiota has been proposed to alleviate neuroinflammation. Recent metagenomics profiling revealed that gut microbiota of KIKO mice shows a decrement of butyrate-producing bacteria and dietary butyrate supplementation improves adipose tissue inflammation (Turchi *et al*., 2023). Dietary butyrate ameliorates microglia-mediated neuroinflammation in several inflammatory mouse models (Jiang et al., 2021; Wei et al., 2023) and improves cognitive decline following neuroinflammatory neurotoxin injection (Ge et al., 2023). In accordance with these data, KIKO mice treated with butyrate show reduced neuroinflammation and improvement of neurobehavioral abilities. In microglial cells downregulating FXN, we observed that butyrate improves the immunometabolic profile via itaconate/Nrf2/GSH pathway. Butyrate shows a strong chemical similarity to β-hydroxybutyrate, a ketone body increased in a FRDA mouse model (Dong et al., 2022). However, comparative analyses revealed that butyrate exerts higher impact in terms of induction of the mitochondrial anti-oxidant genes and inhibition of pro-inflammatory genes (Chriett et al., 2019). It has been suggested that the Nrf2-mediated antioxidant responses induced by butyrate are mediated by Hcar2 (also called as GPR109A) (Guo et al., 2020), which is strongly expressed by CD11b microglial cells (Moutinho *et al*., 2022). Activation of Hcar2 regulates microglial responses to alleviate neurodegeneration in LPS-induced *in vivo* and *in vitro* models (He et al., 2023). In line with this, Hcar2 downregulation restrained the butyrate-mediated anti-inflammatory responses in FXN-deficient microglia.

The current study provides compelling evidence that the loss of FXN is associated with a disruption in mitochondrial activity, rendering microglial cells highly susceptible to inflammatory responses. Furthermore, our research indicates that itaconate plays a pivotal role in mitigating this inflammatory cascade through a Nrf2-mediated mechanism. While the anti-inflammatory properties of butyrate have been extensively documented, our study showcases its remarkable ability to ameliorate the neuroinflammatory phenotype through Hcar2-mediated itaconate/Nrf2/GSH signaling pathway. Remarkably, dietary supplementation of butyrate also demonstrated efficacy in enhancing neuromotor function in a FRDA mouse model. These findings suggest that butyrate holds significant promise as a readily accessible and safe therapeutic option for alleviating FRDA neurological symptoms.

## MATERIALS AND METHODS

### Mice and Treatments

#### WT and KIKO mice

Mouse experimentation was carried out in strict accordance with established standards for the humane care of animals, following approval by the relevant local authorities, including the Institutional Animal Care and Use Committee at Tor Vergata University, and national regulatory bodies (Ministry of Health, licenses no. 324/218-PR and no. 210/202-PR). Both female and male mice were housed in controlled conditions, with a temperature of 21.0°C and a relative humidity of 55.0% ± 5.0%, all while adhering to a 12-hour light/12-hour dark cycle (lights on at 6:00 a.m., lights off at 6:00 p.m.). They were provided with unrestricted access to food and water, and all experimental procedures were conducted in accordance with institutional safety protocols. The female and male Knock-in Knock-out (KIKO) mice were obtained from Jackson Laboratories (#012329), while their female and male littermate C57BL/6 counterparts (WT) were utilized as control subjects. During testing, the researchers were unaware of the genotypes to ensure unbiased results.

The supplementation of butyrate in male mice was performed as previously reported (Turchi *et al*., 2023). Sodium butyrate was incorporated into their food pellets (at a rate of 5 g per kg per day, consistent with their regular daily caloric intake) starting at 4 months of age. This age was selected as it precedes the onset of metabolic changes and continued until the mice reached 8 months of age, which corresponds to a 16-week treatment period. This timeline was chosen because it coincides with the point at which mice typically begin to display metabolic alterations and weight gain. At 8 months of age, the mice were sacrificed by cervical dislocation and cerebellum was immediately processed or stored at -80°C for subsequent analysis.

#### Neurobehavioral Tests

Before rotarod testing, mice were trained for one day on the rotarod at a constant speed of 7 rpm over 1 min, repeated four times. On the day of testing, each mouse was placed on a stationary rod which was then accelerated from 7 rpm to 32 rpm over 5 min. The latency to fall was recorded. This was done over three trials with 60-minute inter-trial intervals.

Before pole testing, mice were acclimated to a wooden pole measuring 30 cm × 1 cm. For the turning time assessment, each mouse was placed head upwards at the top of the pole. The time taken for the mouse to turn 180 degrees downward was recorded. The descent time, from turning to reaching the base of the pole, was subsequently documented.

For tightrope test, a rope measuring 60 cm in length and 1 cm in diameter was securely stretched between two platforms. Each mouse was placed at the center of the rope, and the time taken to reach either platform was noted. This procedure was repeated over three trials with 30-minute intervals between trials.

#### Cells and Treatments

### Primary Microglia Isolation and BV2 cell line

Primary microglia from the cerebellum were isolated following a previously described method (Apolloni et al., 2013). In brief, mice at 5-6 days of age (p6) were euthanized, and the meninges were carefully removed. The cerebellum was then finely chopped and subjected to digestion using 0.01% trypsin and 10 μg/ml DNaseI. After dissociation and filtration through 70 μm filters, cells were suspended in DMEM/F-12 media supplemented with GlutaMAX™ (Gibco, Invitrogen, UK). This media was further supplemented with 10% fetal bovine serum (FBS), 100 Units/ml of gentamicin, and 100 μg/ml of streptomycin/penicillin. The cells were plated at a density of 62,500 cells per cm^2^. After approximately 15 days, a gentle trypsinization was performed using DMEM/F-12 without FBS (0.08% trypsin in DMEM/F-12 without FBS) for 40 minutes at 37°C to eliminate non-microglial cells. The resulting adherent microglial cells, which were highly pure (>98%), were then cultured in a mixture of glial cell-conditioned medium (50%) at 37°C in an atmosphere containing 5% CO2 for 48 hours prior to use.

To isolate cerebella microglia by magnetic cell sorting, cerebellum homogenate as resuspended in 500 mL of magnetic bead buffer (MBB) consisting of PBS without calcium and magnesium, 0.5% w/v bovine serum albumin (BSA), and 2 mM ethylenediaminetetraacetic acid (EDTA). The cell suspension was then filtered through a 30-mm pre-separation filter (Miltenyi, Bergisch Gladbach, Germany) following three filter washes to remove any large particles and debris. The resulting cell suspension was then separated at 300 x g for 5 min at 4°C and resuspended in MBB along with antiCD45 magnetic beads-conjugated antibody (Miltenyi). The cell suspension was incubated for 15 min at 4°C, then diluted with 2 mL of MBB and centrifuged. The resulting cell pellet was resuspended in 500 mL of MBB, applied onto hydrated MS-columns (Miltenyi), washed three times with 500 mL of MBB, and collected with 1 mL of MBB through piston elution. CD45+ cells resuspended in MBB along with antiCD11b magnetic beads-conjugated antibody (Miltenyi). The cell suspension was incubated for 15 min at 4°C, then diluted with MBB and centrifuged. The resulting cell pellet was resuspended in 500 mL of MBB, applied onto hydrated MS-columns (Miltenyi), washed three times with 500 mL of MBB, and collected with 1 mL of MBB through piston elution to obtain CD45+/CD11b+ cells.

Murine BV2 cell line (ATCC) was cultured in DMEM supplemented with 10% FBS and 1% P/S (Life Technologies) and 1% non-essential amino acids (Euroclone). All cells were maintained at 37C in a humidified incubator containing 5% CO2. For gene silencing, BV2 cells were seeded 20000 cell/well. Twenty-four hours after plating, BV2 cells were infected with 25 MOI of FXN shRNA or scramble shRNA (Origene, Rockville, MD, USA) for a total of 500000 viral particles/well. To facilitate viral particle entry in the cells 2ug/mL polybrene (Sigma Aldrich) we added to the culture media. BV2 cells treated with 500 ng/mL lipopolysaccharides (LPS) for 16 h. Sodium butyrate (BUT, 500μM) was added 3 hours before LPS treatment and maintained throughout the experiment. The sodium butyrate concentration was selected based on dose-response experiments conducted on primary adipocytes or bone marrow-derive macrophages stimulated with LPS (500 ng/mL, 16 h). These experiments demonstrated the anti-inflammatory action of the 500 mM concentration while preserving cell viability (Turchi *et al*., 2023).

### Single Cell RNA-sequencing

Single-cell suspensions were prepared for scRNA-seq immediately after cell sorting using the Chromium Single-Cell Reagent Kit from 10x Genomics, following the manufacturer’s protocol. After cell capture and lysis, cDNA was synthesized for each group of captured cells and underwent 12 cycles of amplification. The amplified cDNA from each channel of the Chromium system was utilized to construct an Illumina sequencing library, which was sequenced using the NovaSeq 6000, resulting in approximately 300 million reads per library with a 2×50 read length. Raw reads were aligned to the Mus musculus (mm10) reference genome, and cells were identified using CellRanger count v.7.1.0. Individual samples were combined to create a merged digital expression matrix. The barcodes, features, and matrix files generated by the CellRanger software were used as input for the R program Seurat v4.4.0, (Satija et al., 2015). Low-quality cells were filtered out, retaining only cells with more than 500 features, gene counts greater than 1000, and mitochondrial content less than 10%. Outliers, defined as cells with more than 10,000 features and counts exceeding 20,000, were removed, while retaining genes expressed in at least three cells. This filtering step resulted in 2,400 cells from each sample, which were then merged into a single dataset. Expression levels were normalized using logarithmic transformation. The most variable genes (2,000 features) were selected, and their expressions were scaled across all cells. The dimensionality of the dataset was assessed through Principal Component Analysis (PCA), and the first 20 principal components were used to create a UMAP reduction. Clustering was performed with a resolution parameter set to 0.5. Differentially expressed genes within each cluster were identified using the Wilcoxon rank sum test. Manual cluster labeling was conducted, and annotations were confirmed using the SingleR package with CellDex libraries v1.10.1, (Aran et al., 2019). Enrichment analysis was carried out using ClusterProfileR v4.4.4, PMID: 22455463), which identified the top-5 activated and top-5 suppressed Gene Ontology Biological Processes terms using the gseGO function. Subsequently, the Microglia cluster was isolated from the main dataset and subjected to full reprocessing. Single-cell plots were generated using the GGPlot2 package v3.4.4.

### Bulk RNA-sequencing

Total RNAs were extracted from cells and tissues employing the MiniPrep kit from ZYMO RESEARCH, in adherence to the manufacturer’s instructions. The quantification of total RNA was performed using the Qubit 4.0 fluorimetric Assay from Thermo Fisher Scientific. Libraries were constructed from 50 ng of total RNA through the NEGEDIA Digital mRNA-seq research grade sequencing service provided by Next Generation Diagnostic srl. This service encompassed library preparation, quality assessment, and sequencing on an Illumina NovaSeq 6000 system utilizing a single-end, 75-cycle strategy.The raw data underwent analysis with FastQC v0.12.0 and were subsequently subjected to quality filtering and trimming by Trimmomatic v0.39, (Bolger et al., 2014), employing a Q30 threshold for both leading and trailing ends, with a minimum length of 15 nucleotides. The resultant reads were aligned to the reference genome (mm10) using HISAT2 v2.2.1 (Kim et al., 2019). Quantification of gene expression was accomplished using the featureCounts tool (Liao et al., 2014). Subsequently, the DESeq2 package v1.40.2 (Love et al., 2014) was employed to calculate differentially expressed genes (DEGs) and normalize the expression count matrix. Functional enrichment analysis was carried out using EnrichR or FunRich 3.0 with the Biological Processes Gene Ontology (GO) database.

### Immunophenotyping by Flow Cytometry

Cerebellum was immunophenotyped by high dimensional flow cytometry using a panel containing markers to identify cell types and to assess activation states. The use of these markers allowed us to exclude all cells of no interest based on physical parameters (side and forward scatter) and to gate on specific cells of interest. In particular, cerebellum of WT and KIKO mice was dissociated to single-cell suspension using adult brain dissociation kit from MACS Technology and using GentleMACS (Miltenyi Biotec), according to the manufacturer’s protocol. Cells were first gated on CD45^+^ cells and then on CD45^low^CD11b^+^ to identify microglial cells and to exclude infiltrated macrophages (CD45^high^CD11b^+^) and non-myeloid leukocytes (CD45^high^CD11b^low^). Microglia were further stained for the expression of M1 (anti-CD86 and anti-MHC-II) or M2 (anti-CD206 and anti-Trem2) markers. Samples were acquired on a 13-color Cytoflex (Beckman Coulter) and for each analysis, at least 0.5×10^6^ live cells were acquired by gating on aqua Live/Dead negative cells (Sciarretta et al., 2023).

### Targeted Metabolomics

All data were acquired on a Triple Quad API3500 (AB Sciex) with an HPLC system ExionLC AC System (AB Sciex). For targeted metabolomic analysis, cells were extracted using tissue lyser for 30 sec at maximum speed in 250 µL of ice-cold methanol: water: acetonitrile (55:25:20) containing [U- ^13^C_6_]-glucose 1 ng/µL and [U-^13^C_5_]-glutamine 1ng/µL as internal standards (Merk Life Science, Milan, Italy). Lysates were spun at 15,000 g for 15 min at 4 °C, dried under N_2_ flow at 40 °C and resuspended in 125 µL of ice-cold methanol/water 70:30 for subsequent analyses. Amino acids analysis was performed through the previous derivatization. Briefly, 50 µl of 5% phenyl isothiocyanate in 31.5% ethanol and 31.5% pyridine in water were added to 10 µl of each sample. Mixtures were then incubated with phenyl isothiocyanate solution for 20 min at room temperature, dried under N2 flow, and suspended in 100 µl of 5 mM ammonium acetate in methanol/ H_2_O 1:1. Quantification of different amino acids was performed by using a C18 column (Biocrates, Innsbruck, Austria) maintained at 50 °C. The mobile phases were phase A: 0.2% formic acid in water and phase B: 0.2% formic acid in acetonitrile. The gradient was T_0_: 100% A, T_5.5_: 5% A and T_7_: 100% A with a flow rate of 500µL/min. Measurement of energy metabolites and cofactors was performed by using a cyano-phase LUNA column (50 mm × 4.6 mm, 5 µm; Phenomenex, Bologna, Italy), maintained at 53°C, by a 5 min run in negative ion mode. The mobile phase A was water, while phase B was 2 mM ammonium acetate in MeOH, and the gradient was 50% A and 50% B for the whole analysis, with a flow rate of 500 µL/min. Acylcarnitines quantification was performed using a ZORBAX SB-CN 2.1×150mm, 5µm column (Agilent, Milan, Italy). Samples were analyzed by a 10 min run in positive ion mode. The mobile phases were phase A: 0.2% formic acid in water and phase B: 0.2% formic acid in acetonitrile. The gradient was T_0_: 100% A, T_5.5_: 5% A and T_7_: 100% A with a flow rate of 350µL/min. All metabolites analyzed were previously validated by pure standards, and internal standards were used to check instrument sensitivity. MultiQuant software (version 3.0.3, AB Sciex) was used for data analysis and peak review of chromatograms. Data were normalized on the median of the areas and then used to perform the statistical analysis.

### Lactate Production and Glucose Uptake

Extracellular lactate levels were assessed in the culture medium via an enzyme-based spectrophotometric assay. The procedure involved the collection of cell media, followed by treatment with a 1:2 (v/v) solution of 30% trichloroacetic acid to precipitate proteins. Afterward, the resulting mixture was subjected to centrifugation at 14,000 x g for 20 minutes at 4°C, and the supernatant was carefully collected. Subsequently, the collected supernatant was incubated for 30 minutes at 37°C with a reaction buffer containing glycine, hydrazine, NAD+ (nicotinamide adenine dinucleotide), and LDH (lactate dehydrogenase) enzyme. This incubation allowed for the conversion of lactate to pyruvate, while simultaneously reducing NAD+ to NADH. The concentration of NADH, which is stoichiometrically equivalent to the amount of lactate, was then determined at 340 nm using a spectrophotometer.

To monitor glucose uptake, 2-NBDG probes were used according to manufactures protocols. Flow cytometry analyses were performed by 13-color Cytoflex (Beckman Coulter) and the percentage of 2-NBDG-positive cells was calculated by FlowJo software.

### Quantitative PCR

The total RNA was isolated using TRI Reagent (Sigma-Aldrich). Subsequently, 3 mg of RNA was reverse-transcribed with M-MLV (Promega, Madison, WI). Quantitative PCR (qPCR) was performed in triplicate, using validated qPCR primers confirmed via BLAST searches. The Applied Biosystems Power SYBR Green Master Mix was employed, along with the QuantStudio3 Real-Time PCR System (ThermoFisher, Waltham, MA, USA). mRNA levels were normalized to actin mRNA, and the relative mRNA levels were determined using the 2^-ΔΔCt method. The primers used for reverse transcription quantitative PCR (RT-qPCR) are as follows:

Fxn:orward: 5’-TCTCTTTTGGGGATGGCGTG-3’

Reverse: 5’-GCTTGTTTGGGGTCTGCTTG-3’

Il1b:

Forward: 5’-TGCACCTTTTGACAGTGATG-3’

Reverse: 5’-AAGGTCCACGGGAAAGACAC-3’

Il6:

Forward: 5’-GGATACCACTCCCAACAGA-3’

Reverse: 5’-GCCATTGCACAACTCTTTTCTCA-3’

Rpl8:

Forward: 5’-GGAGCGACACGGCTACATTA-3’

Reverse: 5’-CCGATATTCAGCTGGGCCTT-3’

Nos2:

Forward: 5’-GCCTTCAACACCAAGGTTGTC-3’

Reverse: 5’-ACCACCAGCAGTAGTTGCTC-3’

Hcar2:

Forward: 5’-GAGCAGTTTTGGTTGCGAGG-3’

Reverse: 5’-GGGTGCATCTGGGACTCAAA-3’

Irg1:

Forward: 5’-GCAACATGATGCTCAAGTC-3’

Reverse: 5’-TGCTCCTCCGAATGATACCA-3’

Tnfa:

Forward: 5’-ATGGCCTCCTCATCAGTT C-3’

Reverse: 5’-TTGGTTTGCTACGACGTG-3’

### Immunoblotting

Tissues or cells were lysed in RIPA buffer containing 50 mM Tris-HCl (pH 8.0), 150 mM NaCl, 12 mM deoxycholic acid, 0.5% Nonidet P-40, as well as protease and phosphatase inhibitors. Next, 5 mg of proteins were loaded onto an SDS-PAGE gel and subjected to Western blotting. Nitrocellulose membranes were subsequently incubated with primary antibodies at a 1:1000 dilution. Following this, the membranes were incubated with the appropriate horseradish peroxidase-conjugated secondary antibodies. Immunoreactive bands were detected using a FluorChem FC3 System (Protein-Simple, San Jose, CA, USA) after the membranes were incubated with ECL Prime Western Blotting Reagent (GE Healthcare, Pittsburgh, PA, USA). Densitometric analysis of the immunoreactive bands was performed using the FluorChem FC3 Analysis Software.

### Statistical analysis

The data were presented as the mean ± standard deviation. The specific number of replicates for each dataset is provided in the corresponding figure legend. To evaluate the statistical significance between two groups, a two-tailed unpaired Student’s t-test was conducted. For comparisons involving three or more groups, an analysis of variance (ANOVA) was performed, followed by either Dunnett’s test (for comparisons relative to controls) or Tukey’s test (for multiple comparisons among groups). These statistical analyses were carried out using GraphPad Prism 9 (GraphPad Software Inc., San Diego, CA, USA). In all instances, a significance threshold of p < 0.05 was set.

## ACKNOWLEDGEMENTS

This work was supported by Friedreich’s Ataxia Research Alliance (FARA)-General Research Grant 2021 to D.L.-B. Other supports: Progetto Giovani Ricercatori, Italian Ministry of Health (GR-2018-12367588) to D.L.-B.; FARA-General Research Grant 2020 to KA; Office of the Assistant Secretary of Defense for Health Affairs endorsed by Department of Defense (USA) through the Congressionally Directed Medical Research Programs Award (No. HT9425-23-1-0005) to K.A and D.L.-B.; National Ataxia Foundation (NAF) (821396[RG]) to N.D.A.; #NEXTGENERATIONEU (NGEU) - Ministry of University and Research (MUR), National Recovery and Resilience Plan (NRRP), project MNESYS (PE0000006), A Multiscale integrated approach to the study of the nervous system in health and disease (DN. 1553 11.10.2022) to D.L.-B, K.A and N.D.A. Ministry of University and Research (MUR) Progetto Eccellenza (2023–2027) to the Department of Pharmacological and Biomolecular Sciences “Rodolfo Paoletti” (S.P and N.M.), Università degli Studi di Milano and partially by the Italian Ministry of Health with Ricerca Corrente and 5×1000 funds to NM.

