## Supplemental figures for "Frataxin Deficiency Drives a Shift from Mitochondrial Metabolism to Glucose Catabolism, Triggering an Inflammatory Phenotype in Microglia"

### SUPPLEMENTAL FIGURES AND FIGURE LEGENDS

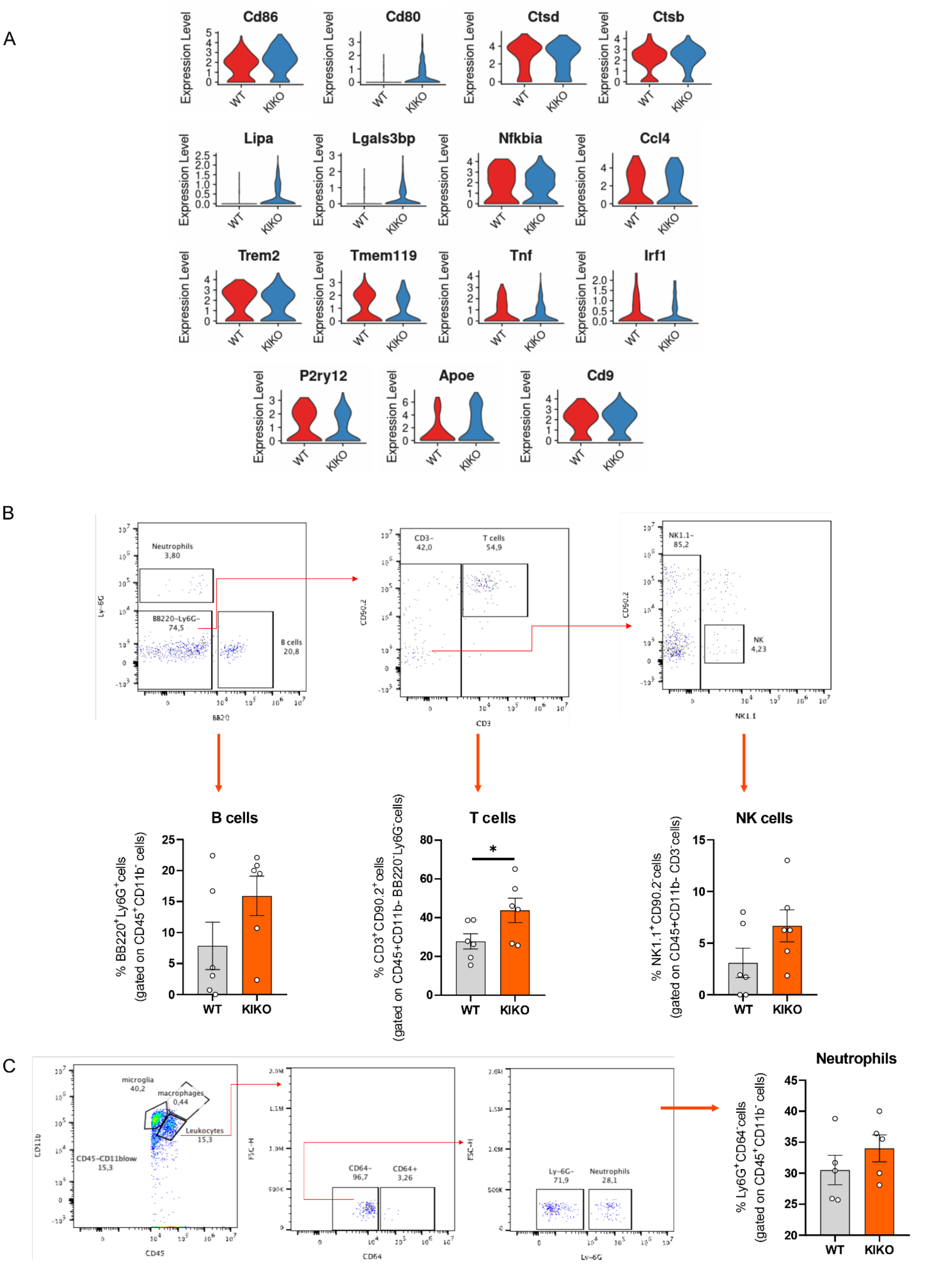


**FIGURE S1. Microglia-derived from cerebellum of KIKO mice show an inflammatory phenotype**. (A) Violin plots reporting inflammatory gene expression in microglia by single cell RNA-seq of cerebellum of WT and KIKO of 6 months old mice (TCF: pool from cerebellum of n=4 mice/group). (**B**) High dimension flow cytometry of T, B and NK cell markers in cerebellum of WT and KIKO of 6 months old mice (n=4/6 mice/group). Data were reported as mean ± SD. Student's t test * p<0.05.

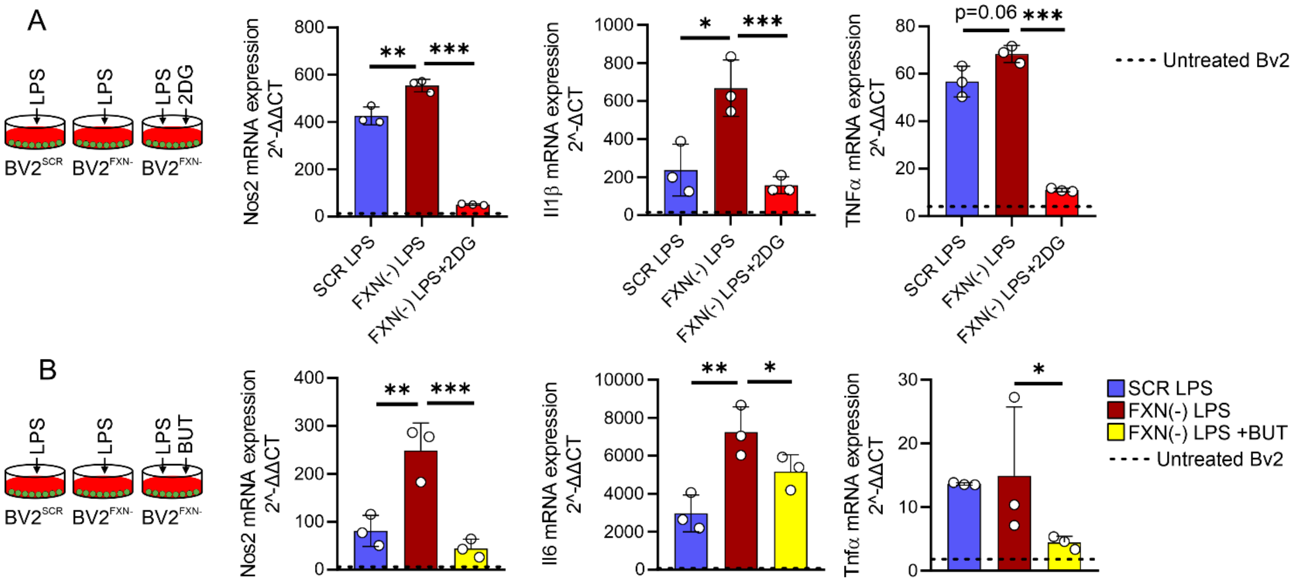


**<u>FIGURE S2</u>. Inhibition of glycolysis or butyrate treatment limit inflammatory response in microglia downregulating frataxin**. (**A-B**) BV2 cells were transfected with lentiviral particles delivering Scr or Fxn sequence and the inflammatory gene expression was analyzed by qPCR. LPS (500 ng/mL for 16 hours) was used to activate BV2 cells. 2-deoxyglucose (2DG, 0.8mM) and sodium butyrate (BUT, 500μM) were added 3 hours before LPS treatment and maintained throughout the experiment. Data were reported as mean ± SD. ANOVA test *p<0.05; p<0.01; *** p<0.001.
